# Spatial transcriptome profiling of *in vitro* 3D tumouroids to study tumour-stroma interactions

**DOI:** 10.1101/2022.12.13.520130

**Authors:** Deniz Bakkalci, Georgina Al-Badri, Wei Yang, Andy Nam, Yan Liang, Syed Ali Khurram, Susan Heavey, Stefano Fedele, Umber Cheema

**Author notes:** Correspondence to: Prof. Umber Cheema, UCL Centre for 3D Models of Health and Disease, Division of Surgery and Interventional Science, University College London, Charles Bell House, 43-45 Foley Street, W1W 7TS, London, UK.

## Abstract

Bioengineering facets of the tumour microenvironment (TME) are essential in 3D tissue models to accurately recapitulate tumour progression. Stromal cells are key components of the TME and their incorporation into 3D biomimetic bioengineered tumour-stroma models is essential to be able to mimic the TME. By engineering tumouroids with distinct tumour and stromal compartments, it has been possible to identify how gene expression is altered by the presence of different stromal cells using spatial transcriptomics. Ameloblastoma is a benign epithelial tumour of the jawbone and in engineered multi-compartment tumouroids increased expression of oncogenes was found where osteoblasts (bone stroma) were present. Engineering a gingival fibroblast stroma resulted in increased matrix remodelling genes in the ameloblastoma tumour. This study provides evidence to show the stromal specific effect on tumour behaviour and illustrates the importance of engineering biologically relevant stroma for engineered tumour models. Our novel results show that an engineered fibroblast stroma causes the upregulation of matrix remodelling genes in ameloblastoma which directly correlates to measured invasion in the model. In contrast the presence of an osteoblast/bone stroma increases the expression of oncogenes by ameloblastoma cells.

## Introduction

The tumour microenvironment (TME) provides essential cues to direct and control tumour progression and provide the basis of key hallmarks of cancer (1). The complexity of the TME derives from both cellular and noncellular components. Cellular components include stromal cells including fibroblasts, immune cells, bone, endothelial cells, and non-cellular components include the matrix components, oxygenation of tissue and cytokines (2). Pertinent extracellular matrix (ECM) composition and stromal cells are required to recreate the TME in engineered tissue models (3). Several different techniques have been studied to mimic cell-cell interactions in the TME both *in vitro* and *in vivo*. Two-dimensional (2D) monolayer co-cultures are widely used as *in vitro* TME models, however they lack many physiological properties, including the tissue architecture. The use of 3D models has made it possible to recapitulate the physiological conditions found in the TME. Multicellular 3D models have been trending due to their ability to conquer the limitations of 2D models (4,5).

One of the challenges of multicellular 3D models is the uncertainty around deciphering which cells signal for specific molecules, in other word, how cells synchronise their signalling and network with other cells (5). Thus, novel strategies are required to unravel this complex communication between tumour and stroma. Rapidly evolving technology in high-plex profiling has led to the development of molecular spatial profiling. The spatial profiling allows cell-type specific characterisation of heterogenous cell populations in the TME. GeoMx Digital Spatial Profiler (DSP) is a platform to spatially resolve biology of the tissue of interest by using digital quantitation of target analytes (6,7). This system utilises barcoded DNA oligos attached to *in situ* hybridisation probes for RNA. The attachment is done by a detection reagent, an ultraviolet (UV)-photocleavable linker (8). The tissue is covered with the detection reagent and the customizable fluorescent morphology markers and then visualised. The regions of interest (ROIs) are used to image the sample following UV light exposure-induced release of the barcoded oligos. These oligos can then be collected by instrument quantitation using nCounter or next generation sequencing (NGS) (8).

The GeoMx DSP works well with small sample size and the process itself is non-destructive, therefore the same section can be profiled multiple times (7). The ROI selection is adjusted based on research of interest. It has been used to study various neoplasms including lung, prostate, breast, or liver cancers and allows for a high level of characterisation of heterogeneous tissue. To our knowledge, this technique has not been tried on any bioengineered in *vitro* samples especially not on odontogenic tumours, but the use of a spatial transcriptomics platform will give insight and demystify the crosstalk between tumour and stroma cells. This study aims to utilise GeoMx DSP to spatially resolve tumour-stroma interaction in *in vitro* 3D tumouroids of ameloblastoma and their native stromal cells.

Ameloblastoma (AM) is a benign odontogenic tumour of the jawbone, which is rare but locally aggressive (9). Ameloblastoma tumour cells interact with the bone and gingival fibroblast stroma, leading to resorption of the surrounding maxillofacial jawbones (10). These interactions regulate the development and progression of the disease, and it is essential to understand the precise mechanisms. So far, our previous studies on 3D ameloblastoma tumouroids have provided novel findings related to disease mechanism (11). Through the careful bioengineering of a connected tumour and stroma compartment, it is possible to study the boundary between these components to profile key pathways in tumour cells which are altered in the presence of and through the interaction with specific stromal cells. The pathways analysed in this study was chose based on our pre-existing gene data (12). Using tumouroid models of ameloblastoma and other cancers, we have already validated invasive marker genes such as matrix metalloproteinases (MMPs) (11,13). Due to the large volume of data generated, this study carefully analyses previous data generated using tumouroid-stroma models to focus on developing clear research questions. These are namely ECM remodelling, invasion and immune-regulation.

This powerful model is fully appreciated when focused regions of interest and cell specific analysis can be conducted at the tumour-stroma interface. By using GeoMx DSP it is possible to analyse key cell populations at this interface and investigate larger cohort of different markers.

## Methods

### Cell Culture

Cell culture conditions were 37°C, 5% CO2, and 21% O2. The immortalised plexiform ameloblastoma cell line, AM-1 was provided by Professor Harada (14). Keratinocyte serum free medium 1X (KSFM) supplemented with KSFM supplements (bovine pituitary extract (BPE) and epidermal growth factor (EGF), human recombinant) was used to culture the AM-1 cells. Primary gingival fibroblasts, Human, Adult (HGF) (PCS-201-018™) were purchased from ATCC and cultured in Dulbecco’s modified Eagle medium (DMEM). Primary human osteoblasts (hOB) from Promocell^®^ (Heidelberg, Germany) were cultured in Promocell^®^ osteoblast growth medium with supplement mix. All media types contained 10% foetal bovine serum (FBS), 100 units/mL penicillin, and 100 μg/mL streptomycin (Gibco™ through Thermo Fisher Scientific, Loughborough, UK).

### 3D Model Fabrication

3D models were engineered using monomeric type I collagen (First Link, Birmingham, UK). and RAFT™ protocol was followed throughout the process. A collagen/cell mix was prepared from 10X Minimal Essential Medium (MEM) (Sigma-Aldrich, Dorset, UK), collagen type I, neutralising agent (N.A) and the cells. N.A was composed of 17% 10 Molar NaOH (Sigma-Aldrich, Dorset, UK) and 83% 10 M HEPES buffer (Gibco™ through Thermo Fisher Scientific, Loughborough). The collagen/cell mix had final volumes of 80% collagen, 10X MEM, 6% N.A. and 4% cells and was kept on ice until it was crosslinked.

The first step in the fabrication of the complex tumouroids was creating the tumour mass of 240 μl of cell/collagen mix with 5×10^4^ AM-1 cells. The mix for the tumour mass was then set into 96-well plates (Corning^®^ Costar^®^, Sigma-Aldrich, Dorset, UK) and incubated in 37°C for 15 min to allow crosslinking. This step was followed by 15 min of plastic compression using RAFT™ absorbers at room temperature (Lonza, Slough, UK). Then the stromal gel mixes containing either no cells/acellular or 1×10^5^ HGFs or 1×10^5^ hOBs were prepared. The first layer, 650 μl of the stromal gel mix was cast on 24-well plate (Corning^®^ Costar ^®^, Sigma-Aldrich, Dorset, UK). The tumour mass was placed in the middle of the first stromal layer and covered by a second stromal layer of 650 μl of the stromal gel mix.

Following crosslinking of the tumouroids for 15 min at 37°C, 24-well RAFT™ absorbers (Lonza, Slough, UK) were used to plastic compress them for 15 min. The gels were supplied with 2 ml of media, which was changed by 50% every 48 hours. The culture period was 14 days.

### Sample Preparation for Spatial Profiler

The protocol from (8) ‘GeoMxTm RNA Assay: High Multiplex, Digital, Spatial Analysis of RNA in FFPE Tissue’ was followed throughout. The sample preparation section was specific to tissue samples. Therefore, this study has established the sample preparation steps for 3D *in vitro* samples.

3D samples were formalin fixed and processed using a processor (Thermo Fisher Scientific, Loughborough, UK) and embedded. The embedded blocks were sectioned into 5 µm sections and these were trimmed appropriately to mount to the VWR Superfrost Plus Slides (Catalogue number 48311-703) (*Figure 1*). The sections were placed within a defined area (36.2 mm long x 14.6 mm wide) in the middle of the slide.

**Figure 1:**
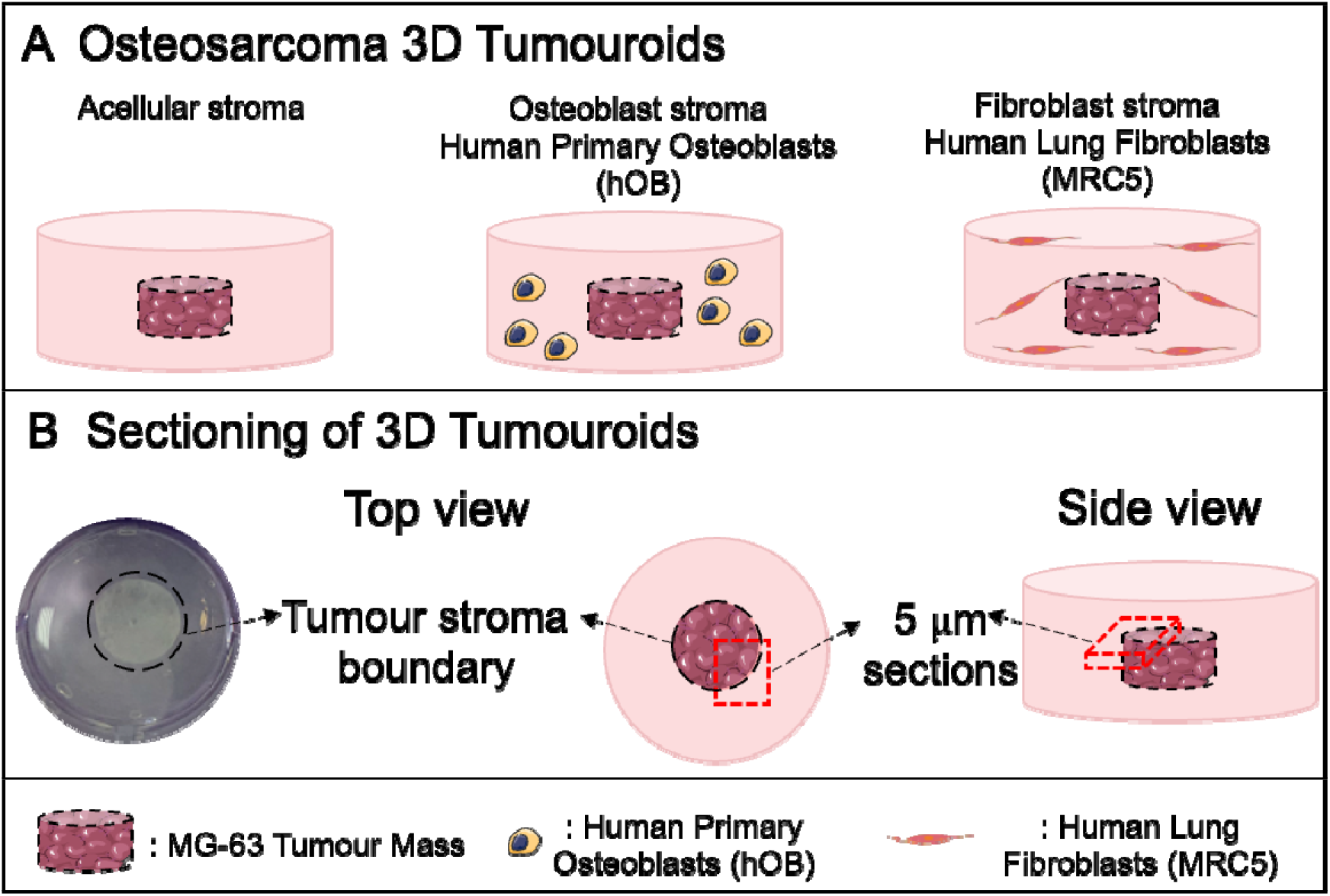
3D model set-up for GeoMx DSP. (A) The 3D tumouroid set-up and (B) sectioning of the 3D tumouroids for GeoMx DSP. Diagrams were created using Smart Servier Medical Art.

### Selection of Region of Interest

Tumour-stroma boundary shown in Figure 1B was chosen as the ROIs for each tissue type. The cellular composition can be segmented into areas of illumination (AOI) by dividing ROIs based on the fluorescent signal of individual morphology marker by filtering the UV-light (15) (*Figure 2A*).

**Figure 2:**
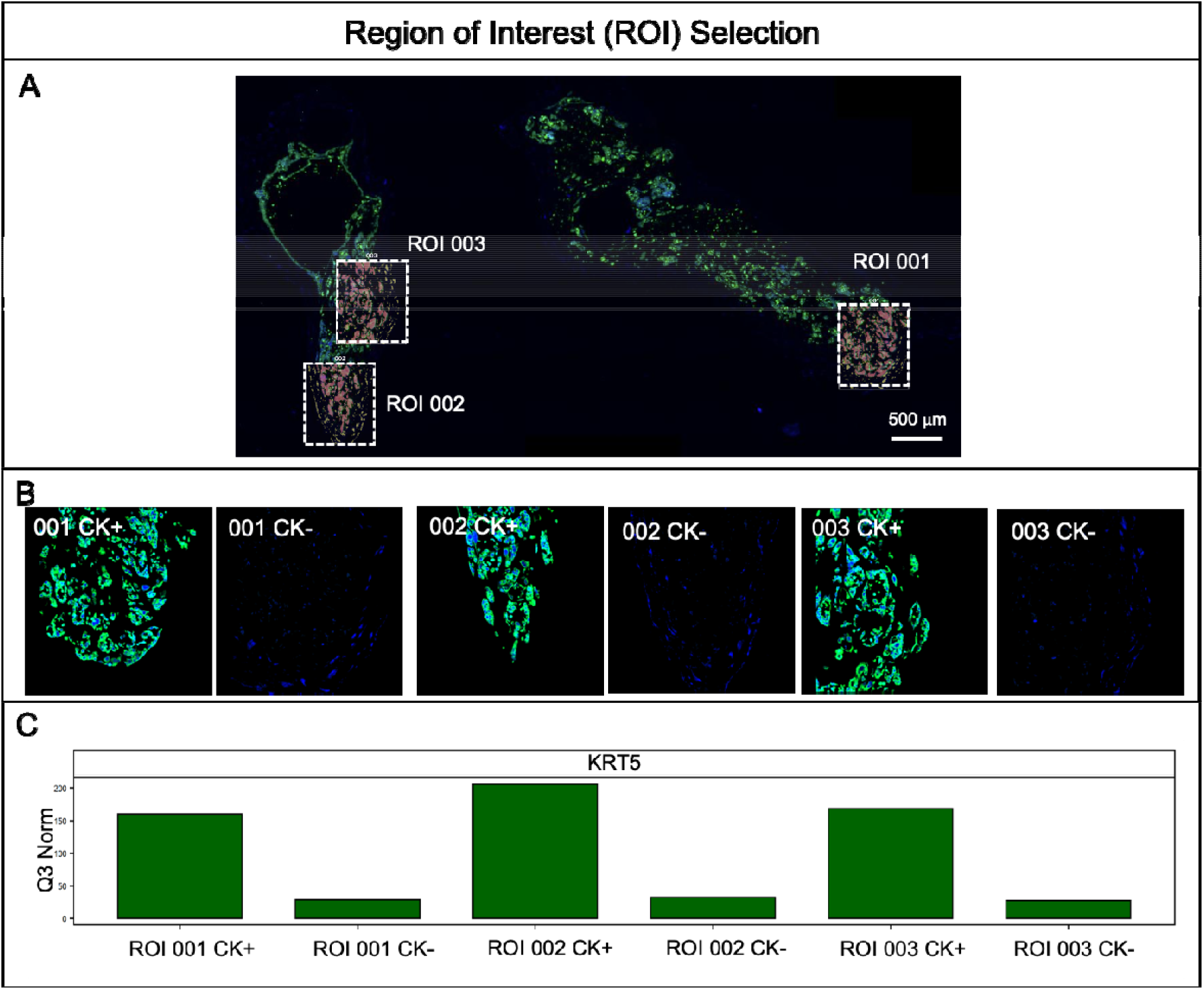
Region of Interest (ROI) selection. (A) Scanned slide of 3D AM-1 tumouroids with HGF stroma and three ROIs, ROI 001, 002, and 003 selected within, scale bar = 500µm. (B) Cytokeratin-positive (CK+) and CK-sections of ROI 001, 002, and 003. (C) Relative mRNA expression of Keratin 5 (KRT5) CK+ and CK-areas following Q3 normalisation.

### NanoString GeoMx Digital Spatial Profiler

The epithelial cell marker pan-cytokeratin (PanCK) was used to identify tumour cells in the samples. PanCK-positive or PanCK-negative cells were profiled individually (15) (*Figure 2 B &C*).

### Histology

Formalin fixed 3D samples were processed overnight using a processor (Thermo Fisher Scientific, Loughborough, UK). The next steps were embedding and sectioning into 5 µm sections. The sections were mounted to Superfrost Plus Slide and the slides were baked at 64°C for 2 hours and then deparaffinised. This was followed by Haematoxylin and eosin (H&E) staining and application of mounting medium for imaging.

### Assay Quality Control and Statistical Analysis

All steps of the assay control and statistical analysis were conducted on GeoMx Digital Spatial Profiler (DSP) Software and Phyton. The sequencing quality is determined based on sufficient saturation and sensitivity of low expressors. Initially raw probe counts were assessed for sequencing quality control (QC) where all of the under-sequenced samples from AOI count analysis were eliminated from the following QCs. The next step was probe QC which is to target mRNAs by multiple probes and the outlier probes were removed. The data was normalised using third quartile (Q3) normalisation. This normalises individual counts to the 75^th^ percentile of signal from their own AOI. The expression levels were presented as counts that quantify RNA level from the readouts of the barcode. The data was evaluated for signal to Limit of Quantitation (LOQ) ratio to test the reliability of the targets. LOQ was calculated as GeoMean(NegProbes) x GeoSD(NegProbes)^2^. Then, the signal from each probe was divided by the LOQ of each AOI.

T-test (non-paired) was used for statistical analysis of gene volcano plots and gene tables (non-paired) with the BH test correction type and tested for by factors. To generate gene volcano plots a custom script for volcano plots by GeoMx Script Hub was used. A p-value of < 0.05 was considered statistically significant and the log_2_ fold change log_2_(FC) value > 0.5 was considered as the notable fold change. The graphs were plotted as log_2_(FC) as x-axis and adjusted p-value as y-axis. Heatmaps were created in Python using hierarchical clustering to visualise statistically significant group gene expression profiles. All statistically significant genes were included in the heatmaps for each pathway unless stated otherwise. Individual genes were plotted using GraphPad Software (La Jolla, CA, USA) and their statistical tests were completed using the GeoMx Digital Spatial Profiler (DSP) Software.

### Selection of genes for pathway panels

All pathway panels were created using set gene lists available in DSP Reactome Target Groups. Invasion pathway gene panel was created from the cell migration genes as well as all MMPs. Matrix remodelling pathway gene panel was composed of ECM proteoglycans, Matrix Remodelling, and ECM Interactions defined in Reactome Target Groups. Immune system pathway gene panel included all Immune System Reactome Target Group.

## Results

### The Ameloblastoma transcriptome is altered in the presence of stromal cells

Tumour-stroma models were engineered using AM-1 cells as a central tumour mass. The central tumour mass was surrounded by specific stromal compartments of dense collagen I containing either no cells, gingival fibroblasts or osteoblasts *(Figure 1A)*. The GeoMx Digital Spatial Profiler (DSP) was used to evaluate the effects on tumour cells of adding specific populations of stromal cells. For the profiler, the area covering the tumour mass-stroma boundary was sectioned *(Figure 1B)* in order to focus and capture tumour cells invading into the surrounding stroma. In each section, 3 ROIs were selected for comprehensive cancer transcriptome analysis *(Figure 2A)*. The ROIs contained both CK+ and CK-cells, and CK+ cells were verified with ∼5-6 timers higher KRT5 expression compared to CK-cells (Figure 2B&C).

The introduction of different physiologically relevant stromal types induced significant changes in the cancer transcriptome of tumour cells. Out of the 1813 transcriptome genes analysed, 1415 genes were above LOQ *(Supplemental Figure 1)*. Volcano plots were generated to overview the changes in the expression profile of AM-1 cells when interacting with different stroma types. Only genes with log_2_ fold-change (log_2_FC) more than 0.5 were further analysed *(Figure 3 A&B&C)*. Initial screening of these volcano plots indicated that the introduction of stromal cells in tumour adjacent compartments induced significant changes (45.3%) in the transcriptome of AM-1 cells compared to acellular stromal compartments. 15.3% of the whole transcriptome was significantly altered when an HGF stroma was added to the AM-1 tumour, 36.7% of genes changed with a hOB stroma, and the comparison between the AM-1 tumour cell expression with either HGF or hOB stroma was 52.4% *(Figure 3)*. The Venn Diagrams in Figure 3D demonstrates number of genes that were significantly downregulated or upregulated with hOB and/or HGF stroma compared to acellular stroma. The number of genes downregulated with hOB stroma was higher than with HGF stroma (Figure 3D).

**Figure 3:**
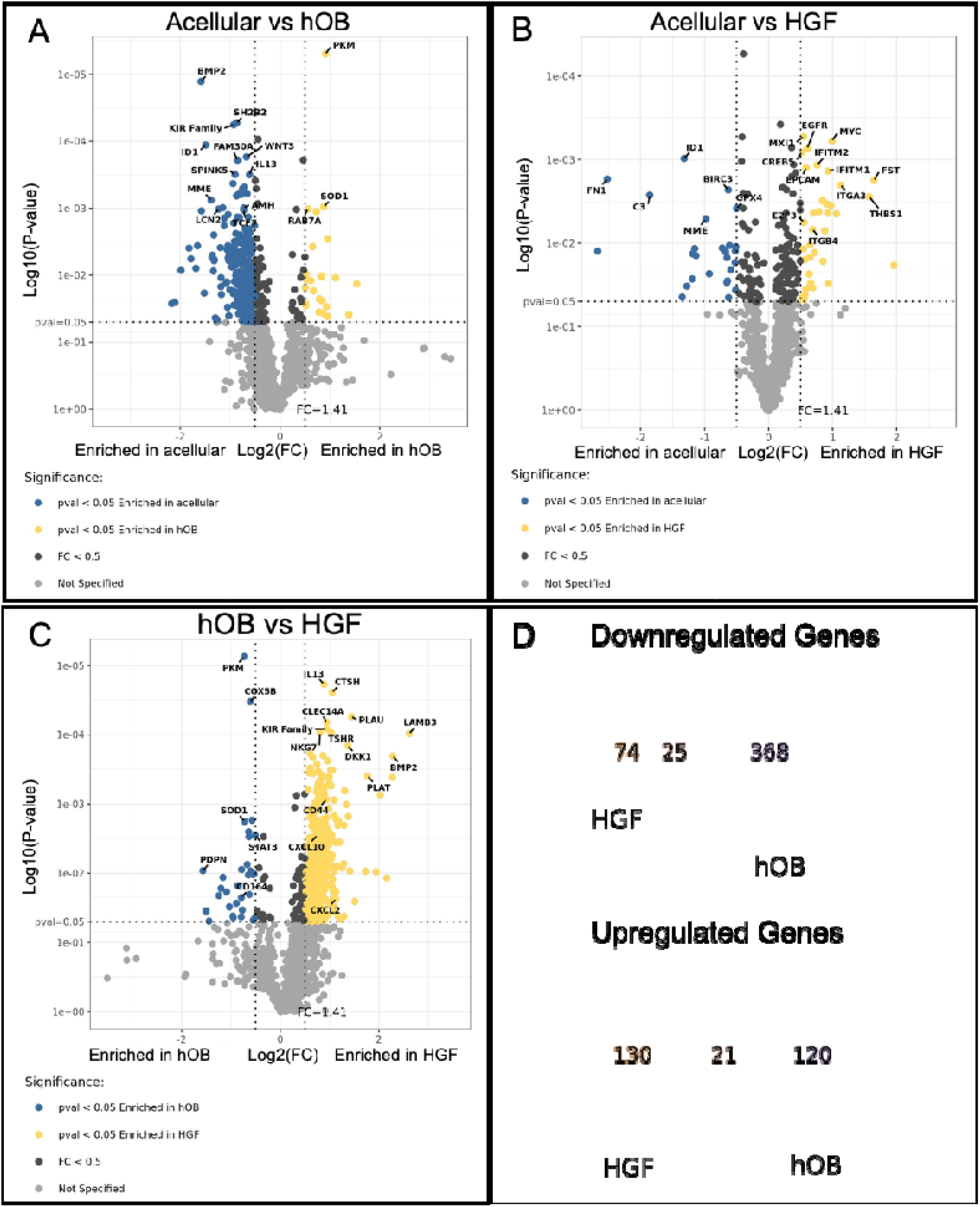
Volcano plots showing Q3 normalised gene counts that are statistically significant, p-value (pval) < 0.05 and fold change (FC) > 0.5 in 3D Ameloblastoma tumouroids at day 14 in CK+ cells. Enrichment of genes were compared among (A) acellular stroma versus (vs) human osteoblast (hOB) stroma, (B) acellular stroma vs human gingival fibroblast (HGF) stroma, and (C) hOB stroma versus HGF stroma. Significance was demonstrated in y-axis as log_10_(P-value) and FC value was presented in x-axis as log_2_(FC). T-test (non-paired), BH test correction, and tested by Factors. (D) Venn Diagrams of number of genes that are significantly upregulated or downregulated in AM-1 cells with HGF stroma and/or hOB stroma compared with acellular stroma.

Considering there were 1415 genes, to better prioritise targets in the case of ameloblastoma, the rest of the analysis was compartmentalised based on different signalling pathways.

### Adding stromal cells cause significant changes in ameloblastoma invasion

Ameloblastoma cells invade into their surrounding stroma (11) within the 3D tumouroid models. Invasion is a tumour specific characteristic driven by different pathways, including epithelial to mesenchymal transition (EMT), migration and cell adhesion pathways (16). Since the sectioned area covered the tumour-stroma boundary, the AM-1 cells that were migrating into the surrounding stroma were spatially profiled.

The invasion of AM-1 cells to their surrounding stroma by day 14 was shown in *Figure 4A*. The invasion data supports our previous finding on the invasion of AM-1 cells to either acellular, HGF and hOB stroma. AM-1 cell invasion was significantly higher into HGF-specific stroma (355±39 µm) compared to acellular stroma (255±66 µm, p<0.05) and to hOB stroma (189±39 µm, p<0.0005) *(Figure 4B)*. The statistically significant change in gene expression with p<0.05 between acellular and hOB was 28.6%, between acellular and HGF was 21.4% and between hOB or HGF versus acellular was 42.9%. These changes suggested that adding stromal cells induced a decrease in specific invasion genes *(Supplemental Table 1) (Figure 4D)*.

**Figure 4:**
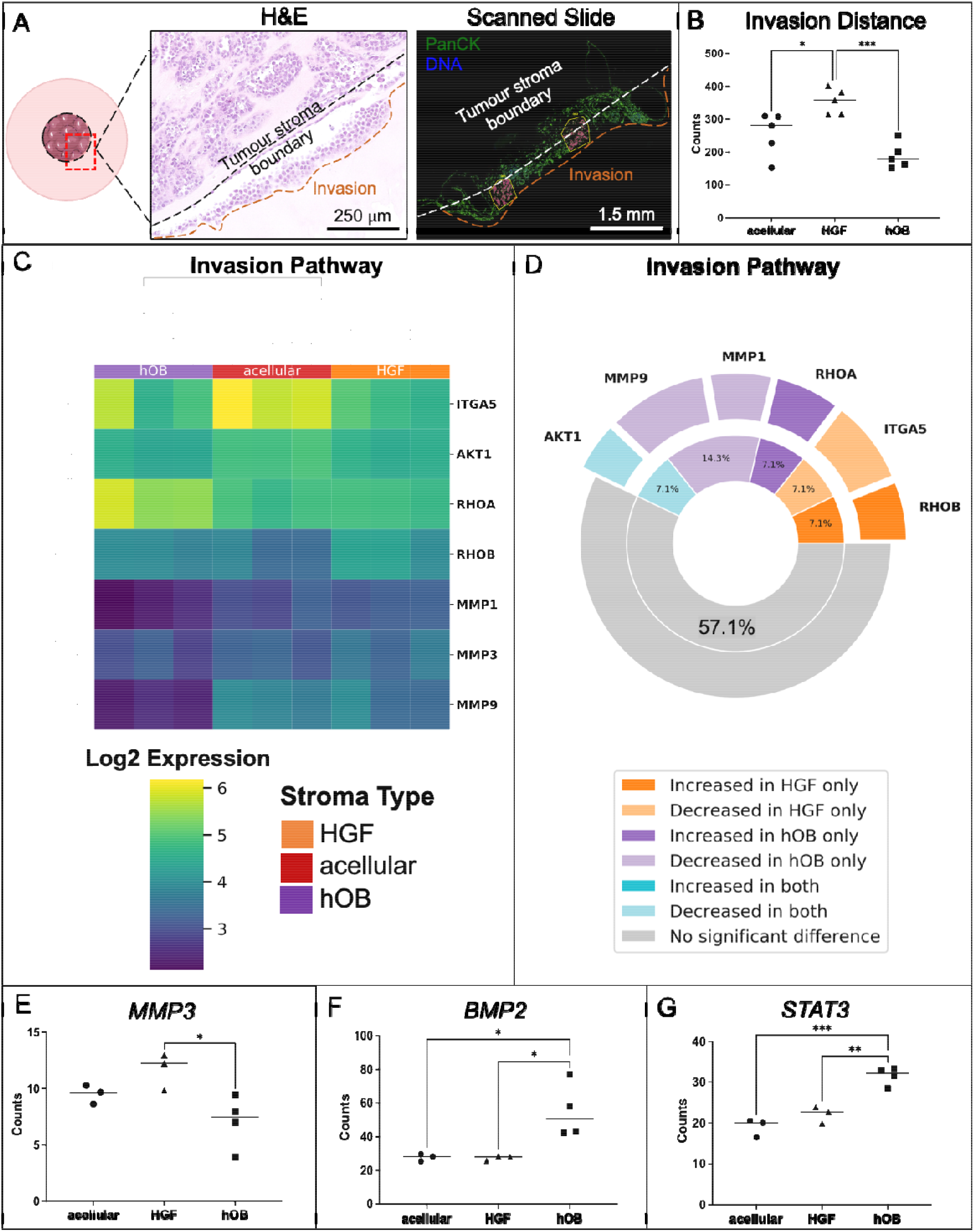
Invasion of AM cells to different stroma. (A) Invasion of AM-1 cells within the 3D tumouorids shown from H&E-stained samples and GeoMx Profiler scanned samples, green = Pan Cytokeratin (PanCK), blue = DNA. scale bars = 250 µm and 1.5 mm respectively. White lines = tumour mass and stroma boundary, orange lines = invasion of tumour cells to the surrounding stroma. (B) Invasion distance of AM-1 cells from the tumour mass to acellular, HGF, and hOB stroma at day 14. (C) **Correlation heatmap of differentially expressed genes in invasion pathway** in AM-1 tumouroids with acellular, HGF, and hOB. Heatmap presents log_2_ change from Mean. T-test (non-paired), BH test correction, and tested by Factors. (D) Plot showing significant gene change. The inner ring represents the percentage of pathway genes significantly changed in the presence of each stroma type. The outer ring indicates the relative fold change in the gene expression observed for each gene in the subgroups. Gene counts of (E) MMP3, (F) BMP2, and (G) STAT3 in AM-1 tumouroids with acellular, HGF and hOB stroma. One-Way ANOVA, Dunnet’s Post Hoc; p-values 0.05 < *, 0.005 < **, and 0.0005 < ***. Diagrams were created using Smart Servier Medical Art.

A heatmap covering EMT, cell migration and MMPs was generated for PanCK+ tissue segments. Different stroma types directly impacted the expression profiles of AM-1 tumour cells. AM-1 tumouroids with hOB stroma demonstrated the greatest change among invasion markers compared to other stroma types *(Figure 4 C&D)*. High expression with hOB stroma varied over different tissue segments. Introducing stromal cells induced an enrichment of certain migration genes but not all. For example, Rho-associated protein kinase (*ROCK1*) was enriched with a hOB stroma. *ROCK1* is important and overexpressed in cell migration and invasion in neoplasms (17). One of the main invasion markers matrix metalloproteinases 3 (MMP3) (18) was ∼2-fold upregulated with HGF stroma compared to hOB stroma in line with the invasion distance data *(Figure 4E)*. The expression of other invasion markers MMP1 and MMP9 were higher with HGF stroma compared to hOB stroma. Interestingly, there was no significant change in some of the other main invasion markers such as *MMP11* and *MMP7* with different stroma types *(Figure 4C)*.

Besides cell migration and adhesion targets, other invasion markers were also assessed. The invasion marker, bone morphogenic protein *(BMP2)* (19) was ∼2-fold upregulated in AM-1 cells cultured with hOB stroma compared to HGF stroma (p<0.05) and acellular stroma (p<0.05) (*Figure 4F*). The invasion and metastasis marker, the signal transducer and activator of transcription 3 *(STAT3)* (20) was significantly upregulated with hOB stroma compared to HGF stroma (p<0.005) *(Figure 4G)*.

### Matrix remodelling ability of the tumour cells is dependent upon their stroma

Tumour cells rely on the interactions with their ECM during cell migration and invasion. There are ∼300 unique matrix macromolecules such as collagens, proteoglycans and glycoproteins such as laminins. Remodelling of the basement membrane composed of collagen IV and laminins is essential for tumour invasion (21). The four main mechanisms of tumourigenic ECM remodelling are ECM deposition, chemical modifications, proteolytic degradation and force-mediated physical remodelling (21). Therefore, the next set of analyses was based on changes related to matrix remodelling targets.

The heatmap for the expression of ECM Proteoglycans targets in AM-1 cells showed varied enrichment levels in particular with the hOB stroma. Certain ECM proteoglycan genes such as the metastatic marker, the amyloid precursor protein *(APP)* (22) significantly upregulated with hOB stroma. AM-1 tumour cells with a HGF stroma presented the greatest enrichment for ECM proteoglycan targets compared to hOB stroma and acellular stroma *(Figure 5A)*. 83.3% of the genes changed by HGF stroma were upregulated, where this percentage was 28.6% with hOB stroma. Overall, 50% of the ECM proteoglycan genes were altered by introducing stromal cells *(Supplemental Table 2)* and *(Figure 6A)*. A similar pattern was observed in ECM interactions targets as well as the matrix remodelling targets *(Figure 5 B&C)*. HGF stroma induced a 57.1% increase among the altered ECM interaction pathway genes in tumour cells and hOB stroma induced a 33.3% increase. The percentage of genes changed in the ECM interaction pathway genes by adding stromal cells was 40.7% *(Supplemental Table 3)* and in the Matrix remodelling pathway was 40.0% *(Supplemental Table 4)*. Adding either hOB and HGF caused significant upregulation of ECM genes involved in tumour progression (23,24) Integrin αvβ6 *(ITGB6)* and *ITGB4* (*Figure 6 A&B*).

**Figure 5:**
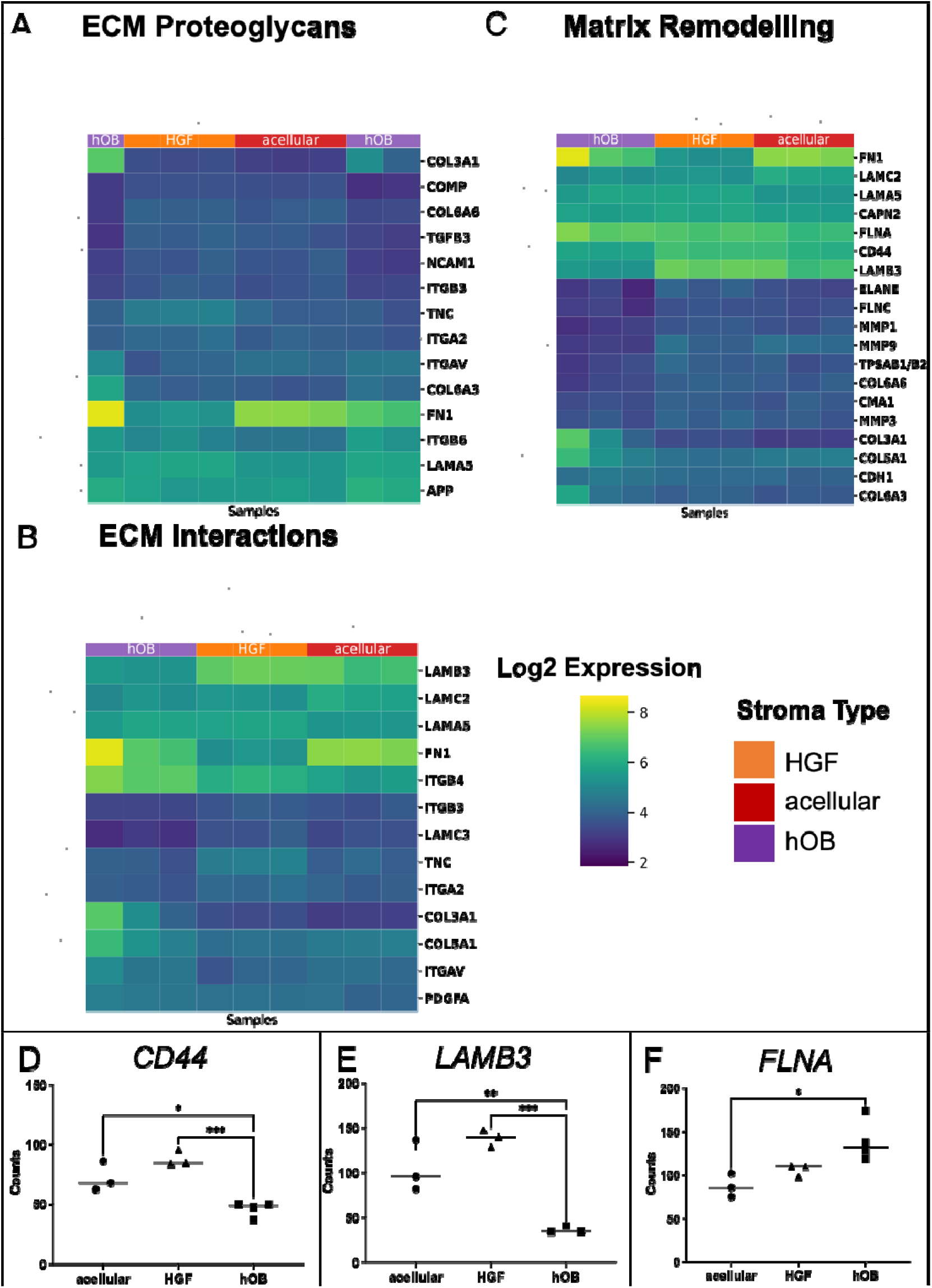
Change in matrix remodelling genes with the introduction of different stromal types. **Correlation heatmap of differentially expressed genes in** (A) ECM proteoglycan pathway, (B) ECM interactions pathway, and (C) matrix remodelling pathway in AM-1 tumouroids with acellular, HGF, and hOB. Heatmap presents log_2_ change from Mean. T-test (non-paired), BH test correction, and tested by Factors. Gene counts of (D) CD44, (E) LAMB3, and (F) COL27A1 in AM-1 tumouroids with acellular, HGF and hOB stroma. One-Way ANOVA, Dunnet’s Post Hoc; p-values 0.05 < *.

**Figure 6:**
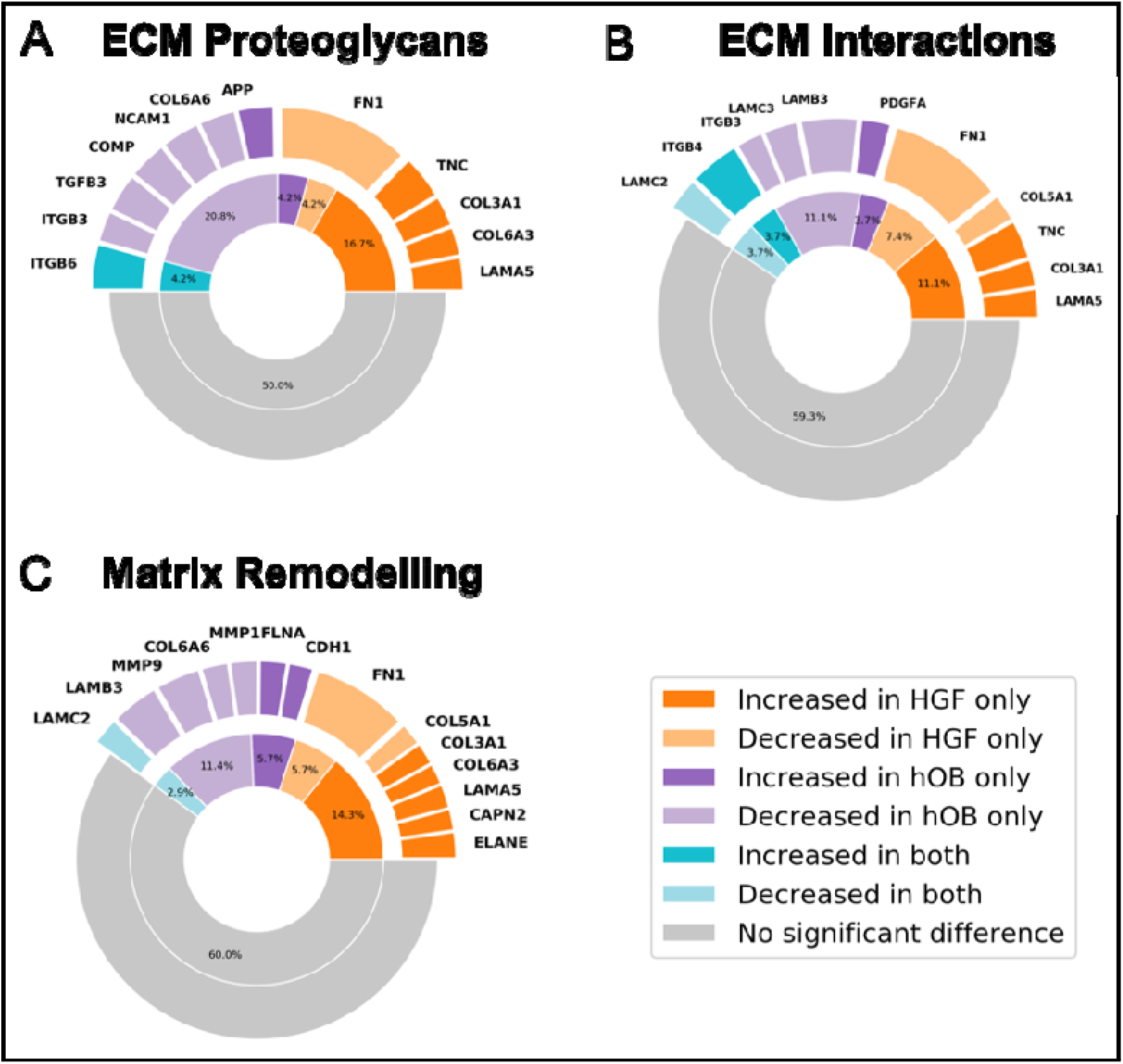
Plots showing significant gene change for (A) ECM Proteoglycans, (B) ECM Interactions, and (C) Matrix Remodelling. The inner ring represents the percentage of pathway genes significantly changed in the presence of each stroma type.

The epithelial mesenchymal transition marker that is highly expressed in primary and metastatic cancers, *CD44* (25), was significantly upregulated where HGF stroma was present compared to hOB stroma (∼2-fold) (p<0.0005) *(Figure 5D)*. The expression of a metastasis marker, the laminin subunit beta-3 *(LAMB3)* was ∼3-fold higher with the HGF stroma compared to hOB stroma (p<0.0005) (Figure 5E). Among matrix remodelling targets, the expression of collagen genes such as collagen type 3 Alpha 2 Chain *(COL3A1)* and *COL6A3* in AM-1 cultured with HGF stroma were higher compared to acellular and hOB stroma *(Figure 6C)*. The matrix remodelling gene Filamin A *(FLNA)* (26), was significantly upregulated with hOB stroma compared to acellular stroma (p<0.05) *(Figure 5F)*.

### Increasing stromal complexity induced enrichment of immune markers by tumour cells

The ECM remodelling by tumour cells influences the inflammatory tumour environment. Components of the ECM act as inflammatory stimuli and drive immune response (21). Therefore, from the cancer transcriptome, the immune pathways have a significant role in understanding how tumour cells communicate with their stroma. The heatmap of the immune system pathway indicate enrichment of the targets by AM-1 cells in hOB stroma and HGF stroma compared to acellular stroma *(Figure 7 A)*.

**Figure 7:**
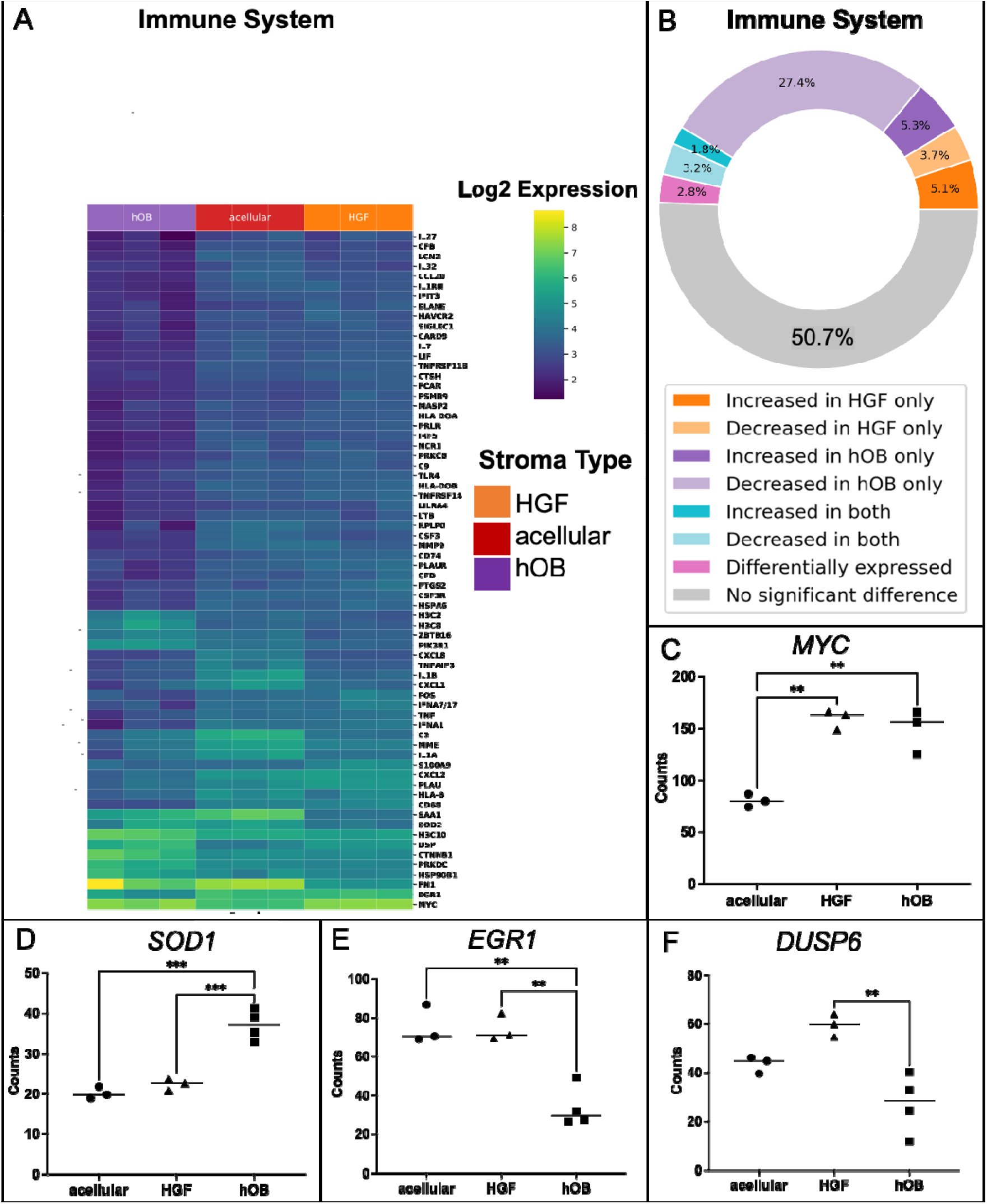
Change in immune system genes with the introduction of different stromal types. **Correlation heatmap of differentially expressed genes** (A) immune system pathway in AM-1 tumouroids with acellular, HGF, and hOB. Data was presented as Log_2_ changes from Mean. T-test (non-paired), BH test correction, and tested by Factors. (B) Plot showing significant gene change. The inner ring represents the percentage of pathway genes significantly changed in the presence of each stroma type. Gene counts of (C) MYC, (D) SOD1, (E) EGR1 and (F) DUSP in AM-1 tumouroids with acellular, HGF and hOB stroma. One-Way ANOVA, Dunnet’s Post Hoc; p-values 0.05 < *, 0.005 < **, and 0.0005 < ***.

Adding either HGF or hOB to the AM-1 stroma induced changes in the expression of 49.3% of genes in the immune system pathway compared to acellular stroma *(Supplemental Table 5)*. In particular HGF stroma induced a significant upregulation in 54.9% of the significantly changed immune system pathway genes compared to acellular stroma. Of the genes that were differentially expressed in the HGF versus the hOB stroma, 85.05% were higher in the HGF stroma. Whereas hOB significantly changed 40.6% of all immune genes, of which 81% were downregulated (*Supplemental Table 5*) and *(Figure 7B)*. The presence of stromal cells resulted in the upregulation of certain oncogenes and downregulated the expression of tumour suppressors, which are part of the immune system pathways. The oncogene, *MYC* (27) was ∼2-fold upregulated in tumour cells with a HGF stroma (p<0.005) and with a hOB stroma (p<0.005) compared to an acellular stroma *(Figure 7C)*. The expression of another oncogene, Superoxide Dismutase 1 *(SOD1)* (28) was significantly higher in AM cells cultured with hOB stroma compared to acellular stroma *(Figure 7D)*. The tumour suppressor markers early growth response 1 *(EGR1) (Figure 7E)* and dual specificity phosphatase 6 *(DUSP6) (Figure 7F)* (29) were downregulated by ∼2-fold in hOB cells compared to the expression in HGF stroma (p<0.005). The expression of EGR1 was also significantly lower with hOB stroma compared to an acellular stroma (p<0.005) *(Figure 7E)*.

## Discussion

Developing 3D models that recapitulate tumour-stroma interactions are essential for modelling the biomimetic tumour microenvironment *in vitro*. 3D tumouroid models allow compartmentalisation where different cell types can be added to distinct compartments to increase model complexity. In these complex tumouroids, mixed cell populations are pooled and then analysed for gene and protein markers (11,30). To date, spatial profiling of the multicompartment 3D models have not been explored. It is difficult to study how the introduction of specific stromal compartment alters or affects tumour cells. This study is the first to capture the area of interest within a bioengineered 3D model and specifically analyse the tumour-stroma boundary.

Bioengineered tumouroids are the first 3D *in vitro* tumour model to be analysed using spatial transcriptomics. Key to this is the sample preparation of spatial transcriptomics which requires embedding and sectioning of the sample. There are limited 3D models that can be processed for histology, including 3D tumouroids and organotypic models (11,31,32). Frozen sectioning is used for spheroid models (33). It is also important to consider the fact that the cell number of 3D models are much lower than normal tissue, and this makes it difficult to section the area of interest. This study described a novel protocol for how to utilise 3D tumouroids for spatial transcriptomics analysis. Within tumouroids, the tumour borders of the tumour mass are visible by eye, thus locating the tumour stroma boundary within the tumouroids is possible during sectioning.

This study assessed changes in the cancer transcriptome of the tumour cells within the 3D tumouroids as the stromal complexity increases. The areas of interests were the tumour mass boundary of ameloblastoma tumouroids to understand the changes in the tumour cells that are invading into the surrounding stroma. The volcano plots generated from the spatial data showed that there are changes in the expression of specific genes in AM-1 tumour cells with different stroma *(Figure 3)*.

To make the story concise, three main pathways from the cancer transcriptome were chosen. Initially, the changes in the invasion pathway targets were compared to the invasion distance data. HGF stroma caused upregulation in invasion markers including *MMP3* (18) and this upregulation was in line with the observed and quantified increase in the invasion distance of AM-1 cells in HGF stroma compared to acellular and hOB stroma. MMPs are well-studied in the case of ameloblastoma, however there is not much data on MMP-3 and most of the reported literature is on MMP-2, -7, and -9 (34,35). Therefore, this study suggests MMP-3 as an invasive marker for ameloblastoma. Although AM-1 cells invaded the shortest distance into a hOB stroma, several invasion genes and metastasis genes such as *RHOA, BMP2* and *STAT3* were upregulated. These markers are associated with later migration compared to MMPs. For example, BMP-2 has been associated with tumour progression in the late stages of gastric cancer (36).

Invasion and matrix remodelling are closely linked (21), therefore it is essential to understand how gene in specific pathways change by adding different stromal cells. Tumour cell-influenced matrix remodelling prepares the TME for tumourigenesis and metastasis. Tumour cells can cause direct or indirect breakdown of the ECM to travel quicker (21). ECM features such as porosity, crosslinking and density also affect cell migration (37).

All these properties were altered with the addition of either hOB or HGF cells in the stroma of AM-1 tumouroids. HGF stroma lead to the greatest enrichment in ECM remodelling targets compared to hOB stroma (Table 4), which is linked to invasion into HGF stroma. Collagen alignment aids the invasion of tumour cells and the collagen chains modulate tumour formation and metastasis (38,39). EMT enhances tumour cell mobility, invasion and metastasis, therefore upregulation of *CD44* in HGF is expected from the invasion distance data *(Figure 5D)*. Our data shows that the hOB stromal cells induce changes in some of the matrix remodelling genes such as *COL27A1*, a gene that is overexpressed by tumour cells and induces ECM production (40). This might be associated with slow migration of the cells compared to HGF and acellular stroma.

Finally, introducing stromal cells induced high enrichment in pathways related to immune system *(Figure 6A)*. Similar enrichment has recently been reported in invasive seminoma germ cell tumours (41). Upregulation of the oncogene *MYC* in the presence of stromal cells, indicated the importance of increasing the complexity of 3D models. Interestingly, the hOB stroma induced downregulation of tumour suppressor genes. It is known that tumour enrichment of tumour markers are linked to enrichment of immune markers (7).

This study highlights how tumour cells are affected and influenced by specific stromal cells within their micro-environment. As tumour cells come into contact with a stromal cell or sense changes in the stroma, significant changes occur. The fact that it is possible to process 3D tumouroids similar to tissue samples with optimised protocols and to be able to conduct spatial analysis, highlight the power of this bioengineered 3D model. The spatial transcriptome data has been used to profile the tumour-stroma boundary within the 3D models. Prior to this work, it was not possible to analyse only CK+ cells or any other cells without disassociating the cells from the tumouroid. This technique allowed us to capture the location of interest, the tumour-stroma boundary. This work will guide future studies that are interested in the applications of spatial transcriptomics in *in vitro* 3D models.

## Limitations

The main limitation is that the GeoMx DSP does not have the same specificity as the single cell sequencing since the platform has 1-10 cells in each spatial spot for analysis. Next steps will include comparing different locations within the tumouroids with each other. The spatial analysis of the whole 3D tumouroid model will enable us to understand the differences in expression patterns within the tumouroid.

## Declarations

### Ethics Approval and Consent to Participate

Not Applicable

### Consent to Publish

Not Applicable

### Availability of data and materials

The authors confirm that data and material in this study are presented in main manuscript.

### Author Contributions

D.B. designed and completed all the 3D experiments and analysed all data except the analysis of the IHC scoring and wrote the manuscript. G.A. helped with pathway data analysis and visualisation in Python. W.Y. helped data analysis using DSP software. A.N. was involved in GeoMx DSP bench work. Y.L. overviewed GeoMx DSP bench work. S.A.K. was involved in conceptualisation, experimental planning, and edited the manuscript. U.C. designed and supervised the project and wrote the manuscript. S.F. supervised the project.

## Acknowledgements

H&E staining was conducted with the guidance of UCL IQPath.

## Abbreviations

AM: Ameloblastoma
AOI: Area of Illumination (AOI)
APP: Amyloid Precursor Protein
BPE: Bovine pituitary extract
*BMP2*: *Bone Moprhogenic Protein 2*
CK: Cytokeratin
*COL3A1*: collagen type 3 Alpha 2 Chain
DMEM: Dulbecco’s modified Eagle medium
DSP: Digital Spatial Profiling
*DUSP6*: Dual specificity phosphatase 6 *(DUSP6)*
ECM: Extracellular matrix
*EGR1*: Early Growth Response
FBS: Foetal bovine serum
HGF: Primary gingival fibroblasts
hOB: Human osteoblasts
IF: Immunofluorescence
*ITGB6*: Integrin αvβ6 (ITGB6)
H&E: Haematoxylin and eosin
*KRT5*: *Keratin 5*
*LAMB3*: laminin subunit beta-3
LOQ: Limit of Quantitation
MEM: Minimal Essential Medium
MIQE: Minimum Information for Publication of Quantitative Real-Time PCR Experiments
MMPs: Matrix metalloproteinases
M: Molar
Min: Minutes
N.A: Neutralising agent
MMP3: matrix metalloproteinases 3
MRC5: Human lung fibroblasts
PanCK: Pan Cytokeratin
*PTHLH*: Parathyroid Hormone Like Hormone
RANK: The tumour necrosis factor (TNF) superfamily members receptor activator of nuclear factor kappa-B receptor
RANKL: The tumour necrosis factor (TNF) superfamily members receptor activator of nuclear factor kappa-B ligand *(TNFSF11)*
ROCK1: Rho-associated protein kinase
ROI: Region of Interest
RT: Room temperature
SD: Standard deviation
SEM: Standard error mean
SOD1: Superoxide Dismutase 1
STAT3: the signal transducer and activator of transcription 3
TGF-β: Transforming growth factor
THBS1: Thrombospondin 1 T
ME: Tumour microenvironment
TNF: Tumour necrosis factor

## References

1. Chen F, Zhuang X, Lin L, Yu P, Wang Y, Shi Y, et al. New horizons in tumor microenvironment biology: challenges and opportunities. BMC Med. 2015;13:45.

2. Chen X, Song E. Turning foes to friends: targeting cancer-associated fibroblasts. Nat Rev Drug Discov. 2019;18(2):99–115.

3. Denton AE, Roberts EW, Fearon DT. Stromal Cells in the Tumor Microenvironment. Adv Exp Med Biol. 2018;1060:99–114.

4. Duval K, Grover H, Han LH, Mou Y, Pegoraro AF, Fredberg J, et al. Modeling Physiological Events in 2D vs. 3D Cell Culture. Physiol. 2017/06/16. 2017;32(4):266–77.

5. Fontana F, Marzagalli M, Sommariva M, Gagliano N, Limonta P. In Vitro 3D Cultures to Model the Tumor Microenvironment. Cancers (Basel). 2021;13(12).

6. Beechem JM. High-Plex Spatially Resolved RNA and Protein Detection Using Digital Spatial Profiling: A Technology Designed for Immuno-oncology Biomarker Discovery and Translational Research. Methods Mol Biol. 2020;2055:563–83.

7. Merritt CR, Ong GT, Church SE, Barker K, Danaher P, Geiss G, et al. Multiplex digital spatial profiling of proteins and RNA in fixed tissue. Nat Biotechnol. 2020;38(5):586–99.

8. Zollinger DR, Lingle SE, Sorg K, Beechem JM, Merritt CR. GeoMx™ RNA Assay: High Multiplex, Digital, Spatial Analysis of RNA in FFPE Tissue BT - In Situ Hybridization Protocols. In: Nielsen BS, Jones J, editors. New York, NY: Springer US; 2020. p. 331–45.

9. Wright JM, Vered M. Update from the 4th Edition of the World Health Organization Classification of Head and Neck Tumours: Odontogenic and Maxillofacial Bone Tumors. Head Neck Pathol. 2017;11(1):68–77.

10. Annegowda VM, Devi HU, Rao K, Smitha T, Sheethal HS, Smitha A. Immunohistochemical study of alpha-smooth muscle actin in odontogenic cysts and tumors. J Oral Maxillofac Pathol. 2018;22(2):188–92.

11. Bakkalci D, Jay A, Rezaei A, Howard CA, Haugen HJ, Pape J, et al. Bioengineering the ameloblastoma tumour to study its effect on bone nodule formation. Sci Rep. 2021;11(1):24088.

12. Bakkalci D, Zaki Abdullah Zubir A, Ali Khurram S, Pape J, Heikinheimo K, Fedele S, et al. Modelling stromal compartments to recapitulate the ameloblastoma tumour microenvironment. Matrix Biol Plus. 2022;100125.

13. Micalet A, Pape J, Bakkalci D, Javanmardi Y, Hall C, Cheema U, et al. Evaluating the Impact of a Biomimetic Mechanical Environment on Cancer Invasion and Matrix Remodelling. Adv Healthc Mater. 2022;e2201749.

14. Harada H, Mitsuyasu T, Nakamura N, Higuchi Y, Toyoshima K, Taniguchi A, et al. Establishment of ameloblastoma cell line, AM-1. J Oral Pathol Med. 1998/07/31. 1998;27(5):207–12.

15. Bergholtz H, Carter JM, Cesano A, Cheang MCU, Church SE, Divakar P, et al. Best Practices for Spatial Profiling for Breast Cancer Research with the GeoMx(®) Digital Spatial Profiler. Vol. 13, Cancers. 2021.

16. Novikov NM, Zolotaryova SY, Gautreau AM, Denisov E V. Mutational drivers of cancer cell migration and invasion. Br J Cancer. 2021;124(1):102–14.

17. Hu C, Zhou H, Liu Y, Huang J, Liu W, Zhang Q, et al. ROCK1 promotes migration and invasion of non✉small✉cell lung cancer cells through the PTEN/PI3K/FAK pathway. Int J Oncol. 2019;55(4):833–44.

18. Liang M, Wang J, Wu C, Wu M, Hu J, Dai J, et al. Targeting matrix metalloproteinase MMP3 greatly enhances oncolytic virus mediated tumor therapy. Transl Oncol. 2021;14(12):101221.

19. Jin H, Pi J, Huang X, Huang F, Shao W, Li S, et al. BMP2 promotes migration and invasion of breast cancer cells via cytoskeletal reorganization and adhesion decrease: an AFM investigation. Appl Microbiol Biotechnol. 2012;93(4):1715–23.

20. Kamran MZ, Patil P, Gude RP. Role of STAT3 in cancer metastasis and translational advances. Biomed Res Int. 2013;2013:421821.

21. Winkler J, Abisoye-Ogunniyan A, Metcalf KJ, Werb Z. Concepts of extracellular matrix remodelling in tumour progression and metastasis. Nat Commun. 2020;11(1):5120.

22. Pandey P, Sliker B, Peters HL, Tuli A, Herskovitz J, Smits K, et al. Amyloid precursor protein and amyloid precursor-like protein 2 in cancer. Oncotarget. 2016;7(15):19430–44.

23. Brendle A, Lei H, Brandt A, Johansson R, Enquist K, Henriksson R, et al. Polymorphisms in predicted microRNA-binding sites in integrin genes and breast cancer: ITGB4 as prognostic marker. Carcinogenesis. 2008;29(7):1394–9.

24. Meecham A, Marshall JF. The ITGB6 gene: its role in experimental and clinical biology. Gene X. 2020;5:100023.

25. Cho SH, Park YS, Kim HJ, Kim CH, Lim SW, Huh JW, et al. CD44 enhances the epithelial-mesenchymal transition in association with colon cancer invasion. Int J Oncol. 2012;41(1):211–8.

26. Zhou J, Wu L, Xu P, Li Y, Ji Z, Kang X. Filamin A Is a Potential Driver of Breast Cancer Metastasis via Regulation of MMP-1. Front Oncol. 2022;12:836126.

27. Casey SC, Baylot V, Felsher DW. The MYC oncogene is a global regulator of the immune response. Blood. 2018;131(18):2007–15.

28. Gomez ML, Shah N, Kenny TC, Jenkins ECJ, Germain D. SOD1 is essential for oncogene-driven mammary tumor formation but dispensable for normal development and proliferation. Oncogene. 2019;38(29):5751–65.

29. James NE, Beffa L, Oliver MT, Borgstadt AD, Emerson JB, Chichester CO, et al. Inhibition of DUSP6 sensitizes ovarian cancer cells to chemotherapeutic agents via regulation of ERK signaling response genes. Oncotarget. 2019;10(36):3315–27.

30. Pape J, Magdeldin T, Stamati K, Nyga A, Loizidou M, Emberton M, et al. Cancer-associated fibroblasts mediate cancer progression and remodel the tumouroid stroma. Br J Cancer. 2020;123(7):1178–90.

31. Pape J, Magdeldin T, Ali M, Walsh C, Lythgoe M, Emberton M, et al. Cancer invasion regulates vascular complexity in a three-dimensional biomimetic model. Eur J Cancer. 2019;119:179–93.

32. Klausner M, Handa Y, Aizawa S. In vitro three-dimensional organotypic culture models of the oral mucosa. Vitr Cell Dev Biol - Anim. 2021;57(2):148–59.

33. Lin C-L, Kuo Y-T, Tsao C-H, Shyong Y-J, Shih S-H, Tu T-Y. Development of an In Vitro 3D Model for Investigating Ligamentum Flavum Hypertrophy. Biol Proced Online. 2020;22(1):20.

34. Kelppe J, Thorén H, Haglund C, Sorsa T, Hagström J. MMP-7, -8, -9, E-cadherin, and beta-catenin expression in 34 ameloblastoma cases. Clin Exp Dent Res. 2021;7(1):63–9.

35. Ohta K, Naruse T, Ishida Y, Shigeishi H, Nakagawa T, Fukui A, et al. TNF-α-induced IL-6 and MMP-9 expression in immortalized ameloblastoma cell line established by hTERT. Oral Dis. 2017;23(2):199–209.

36. Jensen ED, Pham L, Billington Jr CJ, Espe K, Carlson AE, Westendorf JJ, et al. Bone morphogenic protein 2 directly enhances differentiation of murine osteoclast precursors. J Cell Biochem. 2010;109(4):672–82.

37. Pramanik D, Jolly MK, Bhat R. Matrix adhesion and remodeling diversifies modes of cancer invasion across spatial scales. J Theor Biol. 2021;524:110733.

38. Koorman T, Jansen KA, Khalil A, Haughton PD, Visser D, Rätze MAK, et al. Spatial collagen stiffening promotes collective breast cancer cell invasion by reinforcing extracellular matrix alignment. Oncogene. 2022;41(17):2458–69.

39. Huang G, Ge G, Izzi V, Greenspan DS. α3 Chains of type V collagen regulate breast tumour growth via glypican-1. Nat Commun. 2017;8:14351.

40. Arolt C, Meyer M, Hoffmann F, Wagener-Ryczek S, Schwarz D, Nachtsheim L, et al. Expression Profiling of Extracellular Matrix Genes Reveals Global and Entity-Specific Characteristics in Adenoid Cystic, Mucoepidermoid and Salivary Duct Carcinomas. Cancers (Basel). 2020;12(9).

41. Nestler T, Dalvi P, Haidl F, Wittersheim M, von Brandenstein M, Paffenholz P, et al. Transcriptome analysis reveals upregulation of immune response pathways at the invasive tumour front of metastatic seminoma germ cell tumours. Br J Cancer. 2022;126(6):937–47.

