## Supplemental Material for "Spatial transcriptome profiling of *in vitro* 3D tumouroids to study tumour-stroma interactions"

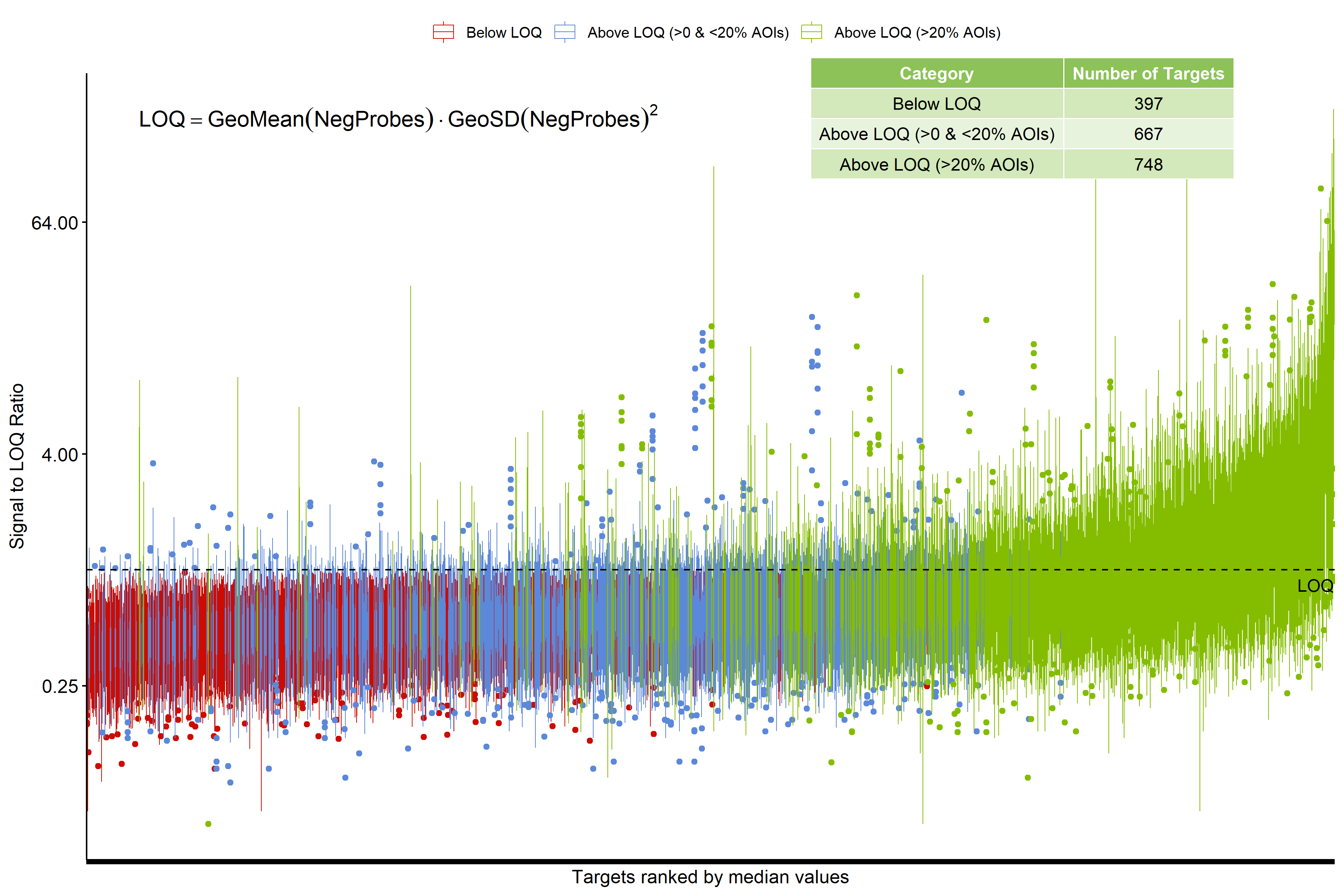

*Supplemental Figure 1: Signal to LOQ Ratio performance. Number of targets/genes that were below LOQ, and above LOQ was demonstrated in the table. LOQ was measured from the equation “GeoMean(NegProbes)xGeoSD(NegProbes)^2^ .*

| **Invasion Pathway**  **Total no. genes: 14** | | | |
| --- | --- | --- | --- |
|  | **% of genes changed** | **% of which increased** | **% of which decreased** |
| HGF vs ac | 21.4 | 33.3 (7.1 of total) | 66.6 (14.3 of total) |
| hOB vs ac | 28.6 | 25 (7.1 of total) | 75 (21.4 of total) |
| HGF OR hOB vs ac | 42.9 |  |  |
|  | **% of genes differentially expressed** | **% of which higher with HGF stroma** | **% of which lower with HGF stroma** |
| HGF vs hOB | 28.6 | 75 (21.4 of total) | 25 (7.1 of total) |
|  | **Genes with increased expression (vs ac)** | **Genes with decreased expression (vs ac)** |  |
| HGF | RHOB | ITGA5, AKT1 |  |
| hOB | RHOA | AKT1, MMP11, MMP9 |  |
|  | **Genes with increased expression (vs hOB)** | **Genes with decreased expression (vs hOB)** |  |
| HGF | MMP1, MMP3, MMP9 | RHOA |  |
| Genes with two significances:   - **AKT1** significantly lower in AM-1 cells cultured with either HGF or hOB stroma compared to acellular stroma. - **MMP9** significantly higher in AM-1 cells cultured with HGF compared to both hOB stroma and acellular stroma. - **RHOA** significantly higher in AM-1 cells cultured with hOB compared to both HGF stroma and acellular stroma. | | | |

*Supplemental Table 1: Invasion pathway and genes that are significantly changed between different stroma types.*

| **ECM Interactions Pathway**  **Total no. genes: 27** | | | |
| --- | --- | --- | --- |
|  | **% of genes changed** | **% of which increased** | **% of which decreased** |
| HGF vs ac | 25.9 | 57.1 (14.8 of total) | 42.9 (11.1 of total) |
| hOB vs ac | 22.2 | 33.3 (7.4 of total) | 66.7 (14.8 of total) |
| HGF OR hOB vs ac | 40.7 |  |  |
|  | **% of genes differentially expressed** | **% of which higher with HGF stroma** | **% of which lower with HGF stroma** |
| HGF vs hOB | 33.3 | 55.6 (18.5 of total) | 44.4 (14.8 of total) |
|  | **Genes with increased expression (vs ac)** | **Genes with decreased expression (vs ac)** |  |
| HGF | LAMA5, ITGB4, COL3A1, TNC | COL5A1, LAMC2, FN1 |  |
| hOB | ITGB4, PDGFA | LAMC2, LAMC3, LAMB3, ITGB3 |  |
|  | **Genes with increased expression (vs hOB)** | **Genes with decreased expression (vs hOB)** |  |
| HGF | LAMB3, ITGA2, LAMC3, ITGB3, TNC | ITGB4, ITGAV, PDGFA, FN1 |  |
| Genes with two significant differences:   - **ITGB4** was significantly increased in AM-1 cells cultured with HGF stroma and hOB stroma (vs acellular stroma), and was significantly higher with hOB stroma compared to HGF stroma. - **LAMC2** was significantly decreased with either HGF or hOB vs acellular stroma. - **ITGB3, LAMC3, and LAMB3** were significantly decreased with hOB compared to HGF and acellular stroma. - **TNC** was significantly increased with HGF stroma compared to hOB and acellular stroma. - **PDGFA** was significantly increased with hOB stroma compared to HGF and acellular stroma. - **FN1** was significantly decreased with HGF stroma compared to acellular and hOB stroma. | | | |

*Supplemental Table 2: ECM interaction Pathway. Table describing gene alteration among different stroma types.*

| **Matrix Remodelling Pathway**  **Total no. genes: 35** | | | |
| --- | --- | --- | --- |
|  | **% of genes changed** | **% of which increased** | **% of which decreased** |
| HGF vs ac | 22.9 | 33.3 (7.1 of total) | 66.6 (14.3 of total) |
| hOB vs ac | 20 | 25 (7.1 of total) | 75 (21.4 of total) |
| HGF OR hOB vs ac | 40.0 |  |  |
|  | **% of genes differentially expressed** | **% of which higher with HGF stroma** | **% of which lower with HGF stroma** |
| HGF vs hOB | 37.1 | 84.6 (31.4 of total) | 15.4 (5.7 of total) |
|  | **Genes with increased expression (vs ac)** | **Genes with decreased expression (vs ac)** |  |
| HGF | ELANE, CAPN2, LAMA5, COL6A3, COL3A1 | COL5A1, LAMC2, FN1 |  |
| hOB | CDH1, FLNA | MMP1, COL6A6, MMP9, LAMB3, LAMC2 |  |
|  | **Genes with increased expression (vs hOB)** | **Genes with decreased expression (vs hOB)** |  |
| HGF | ELANE, MMP1, CAPN2, COL6A6, TPSAB1/B2, MMP9, CMA1, MMP3, LAMB3, CD44, FLNC | CDH1, FN1 |  |
| Genes with two significances:   - **LAMC2** significantly lower in AM-1 cells cultured with HGF stroma and hOB stroma compared to acellular stroma. - **ELANE** and **CAPN2** significantly higher with HGF compared to both hOB and ac - **MMP1, COL6A6, MMP9, LAMB3** all significantly lower with hOB compared to HGF and ac - **CDH1** significantly higher with hOB compared to HGF and ac - **FN1** significantly lower with HGF compared to hOB and ac | | | |

*Supplemental Table 3: Matrix Remodelling Pathway. Genes that are significantly changed between different stroma types.*

| **ECM Proteoglycans**  **Total no. genes: 24** | | | |
| --- | --- | --- | --- |
|  | **% of genes changed** | **% of which increased** | **% of which decreased** |
| HGF vs ac | 25.0 | 83.3 (20.8 of total) | 16.7 (4.7 of total) |
| hOB vs ac | 29.2 | 28.6 (8.3 of total) | 71.4 (20.8 of total) |
| HGF OR hOB vs ac | 50.0 |  |  |
|  | **% of genes differentially expressed** | **% of which higher with HGF stroma** | **% of which lower with HGF stroma** |
| HGF vs hOB | 45.8 | 63.6 (29.2 of total) | 36.4 (16.7 of total) |
|  | **Genes with increased expression (vs ac)** | **Genes with decreased expression (vs ac)** |  |
| HGF | LAMA5, ITGB6, COL6A3, COL3A1, TNC | FN1 |  |
| hOB | APP, ITGB6 | COL6A6, NCAM1, COMP, TGFB3, ITGB3 |  |
|  | **Genes with increased expression (vs hOB)** | **Genes with decreased expression (vs hOB)** |  |
| HGF | COL6A6, NCAM1, COMP, TGFB3, ITGA2, ITGB3, TNC | APP, ITGB6, ITGAV, FN1 |  |
| Genes with two significances:   - **ITGB6** significantly higher in AM-1 cells cultured with HGF stroma and hOB stroma compared to acellular stroma, also significantly higher with hOB stroma compared to HGF (ac < HGF < hOB) - **COL6A6, NCAM1, COMP, TGFB3, ITGB3** were all significantly lower with hOB stroma compared to both HGF stroma and acellular stroma. - **TNC** significantly higher with HGF stroma compared to both hOB stroma and acellular stroma. - **APP** significantly higher with hOB stroma compared to HGF stroma and acellular stroma. - **FN1** significantly lower with HGF stroma compared to both hOB stroma and acellular stroma. | | | |

*Supplemental Table 4: ECM Proteoglycans Pathway. Genes that are significantly changed between different stroma types.*

| **Immune Pathway**  **Total no. genes: 493** | | | |
| --- | --- | --- | --- |
|  | **% of genes changed** | **% of which increased** | **% of which decreased** |
| HGF vs ac | 16.6 | 54.9 (9.1 of total) | 45.1 (7.5 of total) |
| hOB vs ac | 40.6 | 19.0 (7.7 of total) | 81 (32.9 of total) |
| HGF or hOB vs ac | 49.3 |  |  |
|  | **% of genes differentially expressed** | **% of which higher with HGF** | **% of which lower with HGF** |
| HGF vs hOB | 55.6 | 85.0 (47.3 of total) | 15.0 (8.3 of total) |
| Number of significant genes: 331 | | | |

*Supplemental Table 5: Immune Pathway. Table representing % of genes that are changed among different stroma.*
